# eRREMS expands regulatory CpG coverage in reduced representation methylome sequencing

**DOI:** 10.64898/2026.08.03.740891

**Authors:** Uxue Lazcano, Monika Gonzalez-Lopez, Nuria Macías-Cámara, Laura Barcena, Ianire Astobiza, Antonio Rosino, Julian Tudela, Enrique Gonzalez-Billalabeitia, Arkaitz Carracedo, Ana M. Aransay, Isabel Mendizabal

## Abstract

Profiling the regulatory DNA methylation landscape remains technically challenging. Reduced-representation bisulfite sequencing (RRBS) misses distal enhancers and non-canonical regulatory elements, while whole-genome bisulfite sequencing (WGBS) distributes reads genome-wide, limiting consistent recovery of informative CpGs across sample collections. Here we present eRREMS (enhanced Reduced Representation Enzymatic Methylation Sequencing), addressing both limitations by combining MspI and HaeIII digestion with enzymatic cytosine conversion. eRREMS approximately doubles regulatory CpG recovery relative to standard RRBS, outperforming TaqαI-based extended protocols, with gains concentrated in dynamically regulated enhancers, alternative promoters, and splicing-associated regions. HaeIII-specific CpGs capture additional complex trait heritability beyond standard RRBS. At modest, comparable sequencing depths, eRREMS achieves greater cohort-level completeness for regulatory and dynamically variable CpGs than WGBS and requires substantially fewer reads. CpG capture is stable across enzymatic conversion kits and library preparation conditions, supporting broad adoption. eRREMS provides a cost-efficient and reproducible strategy for scalable methylome profiling of regulatory CpGs across cohorts.

## Introduction

Cytosine methylation contributes to the stability and transcriptional regulation of mammalian genomes, with established roles in cell identity, developmental programs, and disease-associated transcriptional rewiring^1,2^. Early studies focused primarily on promoter-proximal CpG islands, but regulatory DNA methylation extends into distal enhancers, alternative promoters, and intragenic elements^3–6^, where it accounts for a significant fraction of the heritability of complex traits^7^. These non-canonical elements frequently exhibit tissue-specific and developmentally dynamic methylation patterns and have been linked to aging, cell type identity and cell-state transitions^2,8–10^. Comprehensive and reproducible profiling of DNA methylation across these regulatory regions therefore remains a key objective in epigenomic research.

DNA methylation profiling strategies face a fundamental tradeoff between CpG coverage breadth and per-site depth. Whole genome bisulfite sequencing (WGBS)^11,12^, considered the gold-standard, provides single-base resolution across ∼28 million CpGs genome-wide, but achieving high per-CpG sequencing depth across the entire genome is costly. More cost-effective approaches such as Reduced Representation Bisulfite Sequencing (RRBS)^13^ enrich for CpG-dense regions through MspI restriction enzyme digestion. However, the preference for promoter-proximal CpG islands^14^ leaves the distal and intragenic regulatory elements poorly represented. This tradeoff is most consequential at cohort scale, where the relevant question is not how many CpGs are theoretically accessible, but how many can be quantified reproducibly across individuals at sufficient depth for downstream analysis. As epigenomic studies scale toward biobank-sized cohorts^15^, the cost of achieving high and uniform CpG coverage across large sample collections risks leaving methylome profiling behind other omics modalities, while array-based alternatives sacrifice coverage breadth and introduce platform-specific technical limitations^16^.

Prior reduced representation protocols have explored alternative enzyme combinations to extend CpG recovery beyond MspI-based RRBS^17^, an approach termed enhanced RRBS (eRRBS), typically through TaqαI addition^18^. Separately, extended fragment recovery around MspI restriction sites (extended-representation bisulfite sequencing, XRBS) has been used to expand regulatory element coverage in low-input and single-cell contexts^19^. Incorporation of HaeIII in RRBS workflows improves coverage of distal regulatory elements in low-input and single-cell applications^20–22^. More recently, enzymatic cytosine conversion, a three-step oxidation-deamination process^23^, has been integrated into reduced representation approaches^24^ to mitigate the DNA damage associated with bisulfite treatment^25,26^. However, a systematic evaluation of HaeIII in combination with enzymatic conversion focusing on regulatory CpG coverage, biological relevance, and cohort-level completeness under realistic sequencing budgets has not been undertaken.

Here we present eRREMS (enhanced Reduced Representation Enzymatic Methylation Sequencing), a reduced representation strategy combining MspI and HaeIII digestion with enzymatic cytosine conversion to extend methylome coverage into non-canonical regulatory elements. We benchmark this approach against classical RRBS in matched biological samples and against WGBS in an independent deeply sequenced cohort. eRREMS substantially expands access to the distal and intragenic regulatory methylome beyond standard RRBS, including elements that disproportionately harbor complex trait heritability. Compared to WGBS, it achieves greater cohort-level completeness at lower read depths, while maintaining methylation concordance across library preparation protocols. Together, these properties position eRREMS as a practical alternative to existing approaches for large-scale regulatory epigenomics.

## Results

### Restriction enzyme selection and library design

Enzyme selection was informed by prior comprehensive *in silico* evaluation of 4-base cutters, which identified HaeIII (GG|CC) as a strong complement to MspI (C|CGG) owing to broader representation across CpG-poor genomic contexts^27^. TaqαI (T|CGA) was included to benchmark against the most widely adopted MspI complement in current extended RRBS protocols^18^. To determine the optimal digestion strategy, we performed *in silico* digestion of the human genome comparing joint versus independent enzyme application. Independent digestion followed by pooling yielded approximately 11.6% more fragments within the 100-500 bp size selection window than joint digestion for the MspI + HaeIII combination (4.5 million vs 4 million; **Supplementary Table 1**). Each enzyme was therefore applied in independent digestion reactions, with products combined after size selection for all subsequent analyses.

To enable direct comparison of restriction enzyme combinations and conversion chemistries, all libraries were generated from the same digested DNA material (**Fig. 1**). Genomic DNA from three fresh-frozen localized prostate cancer samples was independently digested with MspI, TaqαI, or HaeIII (**Supplementary Fig. 1a**). Fragments were size-selected for 100-500 bp inserts (**Supplementary Fig. 1b**). Equal amounts (5 ng) from each digestion were combined to produce three configurations per sample. For MspI-only libraries, two equal aliquots of MspI-digested DNA (5 ng each, 10 ng total) were combined to match the total input of dual-enzyme configurations: MspI-only (M), MspI + TaqαI (MT), and MspI + HaeIII (MH; see Methods and **Supplementary Note** for detailed protocol). We processed each configuration in parallel with bisulfite conversion^28^ and enzymatic conversion^23^, yielding 18 libraries in total (**Supplementary Fig. 1c)**. Spike-in controls, methylated pUC19 and unmethylated λ-bacteriophage DNA, were included to assess conversion efficiency (**Supplementary Table 2**). All libraries were sequenced simultaneously using 150 bp paired-end reads and processed through a unified computational pipeline.

**Figure 1.**
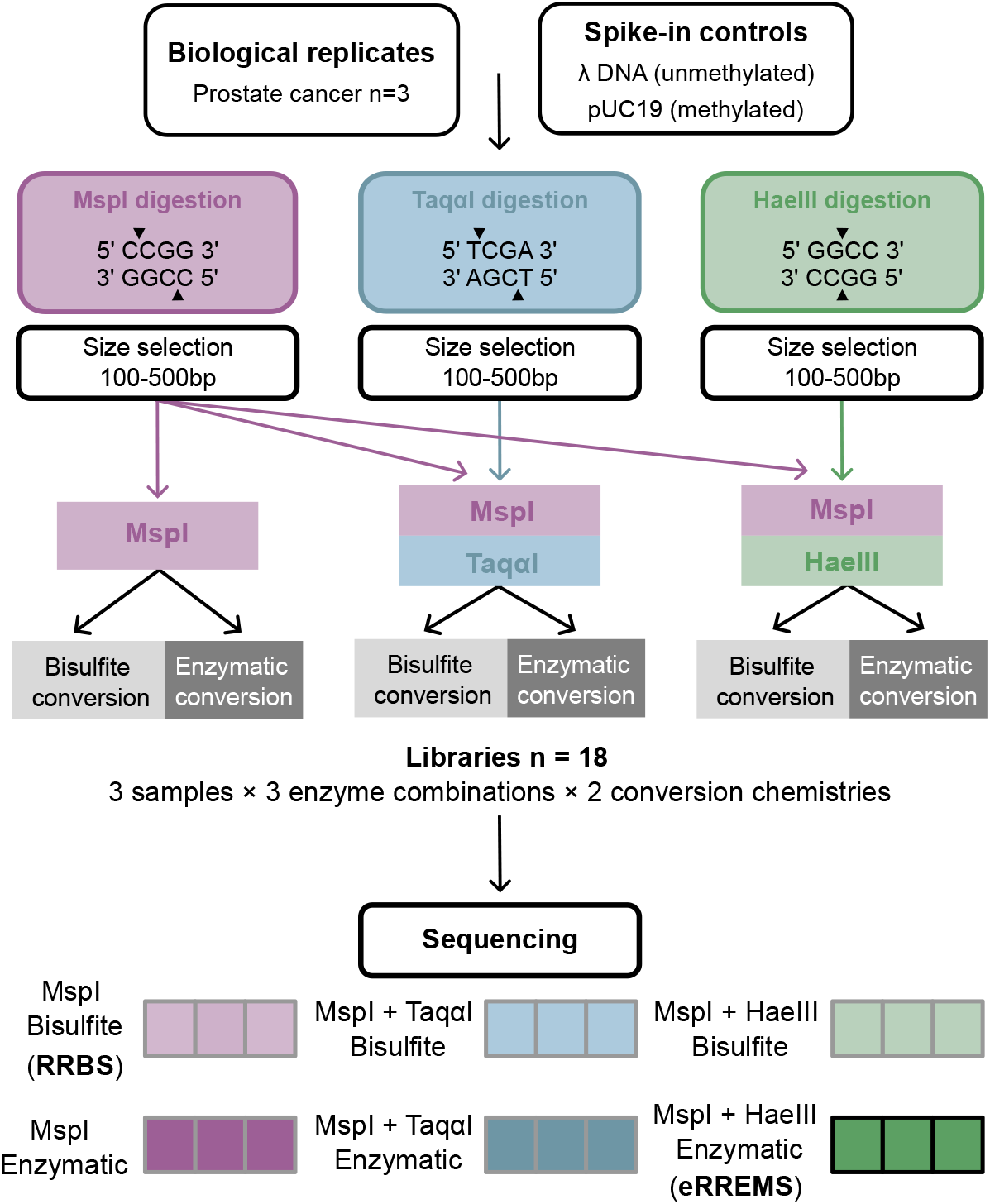
**Study design for systematic benchmarking of restriction enzyme combinations and conversion chemistries**. Schematic outline of the study design including the selection of biological samples, DNA methylation conversion controls, enzymatic digestions, conversion methods, library preparation and sequencing. The MspI + HaeIII enzymatic conversion configuration, constituting eRREMS (enhanced Reduced Representation Enzymatic Methylation Sequencing), is highlighted in the library color scheme; remaining configurations serve as benchmarks, including RRBS (Reduced Representation Bisulfite Sequencing).

The MH enzymatic configuration is hereafter referred to as eRREMS (enhanced Reduced Representation Enzymatic Methylation Sequencing), the method evaluated in this study; MT and M configurations were included as benchmarks. For clarity, libraries are labeled by their configuration and conversion chemistry throughout comparative figures.

#### Restriction enzyme choice and conversion chemistry jointly shape fragment properties and sequencing yield

Enzymatic conversion libraries showed substantially higher yields than bisulfite-converted counterparts prior to sequencing (pooled mean 5,226 ng vs 1,984 ng; **Fig. 2a**) with correspondingly higher raw read counts after sequencing (pooled mean 95 million vs 75 million paired-end reads; **Supplementary Fig. 1d** and **Supplementary Table 3**), consistent with bisulfite-induced DNA strand breaks reducing library yield^25,29^. After mapping, enzymatic libraries exhibited longer insert sizes (pooled mean 281 vs 228 bp; *P* < 2.2x10^-16^, Kolmogorov-Smirnov test; **Fig. 2b**), higher insert GC content (mean 0.49 vs 0.44; *P* < 2.2x10^-16^, Kolmogorov-Smirnov test; **Fig. 2c**), consistent with bisulfite-induced fragmentation and preferential degradation of cytosine-rich DNA^23,25,26^. Because all libraries derived from the same restriction digests and mapping rates were comparable (**Supplementary Table 3**), these differences are attributable to conversion chemistry rather than digestion bias or alignment efficiency.

**Figure 2.**
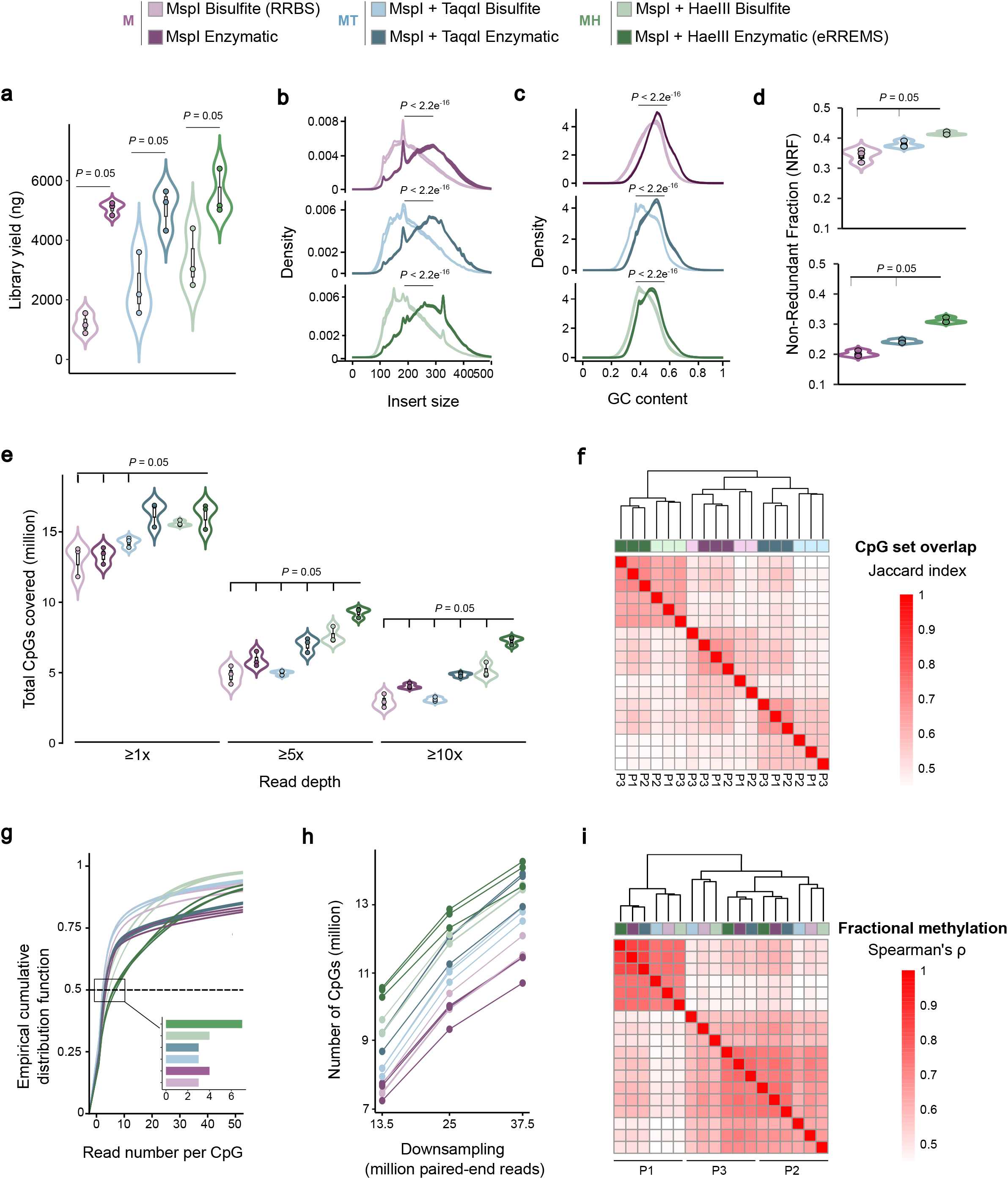
Library characteristics and sequencing performance. **a.** Library yield (ng) prior to sequencing. **b**. Inserts size distributions (base pairs) across libraries. **c**. GC content distributions of mapped inserts. **d**. Library complexity measured as non-redundant fraction. **e.** Number of CpG sites detected at ≥1x, ≥5x and ≥10x coverage across libraries. **a-e.** Significance was obtained with one-tailed Wilcoxon signed-rank test comparing MH enzymatic with each of the other libraries. Only significant comparisons (*P* = 0.05) are indicated. *P* = 0.05 represents the minimum achievable value for the sample size used (n=3 per group). **f**. Pairwise Jaccard similarity indices for CpG sites detected at ≥5x coverage across all libraries. Each library is color-labeled by its configuration (M, MT, or MH; bisulfite or enzymatic conversion) and individual ID (P1–P3). Libraries were hierarchically clustered based on Jaccard similarity values. **g**. Empirical cumulative distribution function of CpGs read depth across libraries. Inset shows median read depth per CpG (50^th^ percentile) for each library configuration. **h**. CpG recovery as a function of sequencing depth after downsampling. **i**. Heatmap of pairwise Spearman correlations of fractional DNA methylation across libraries. Correlations were computed using CpGs covered at >=5x in both samples, with fractional methylation defined as C/(C+T). Only shared CpGs across all samples were considered. Each library is color-labeled by its configuration (M, MT, or MH; bisulfite or enzymatic conversion) and individual ID (P1–P3). Libraries were hierarchically clustered based on correlation values.

Library complexity, measured as non-redundant fraction (NRF), revealed distinct effects of conversion method and enzyme choice. NRF was higher in bisulfite-converted libraries (pooled mean 0.38) relative to enzymatic libraries (pooled mean 0.25; *P* = 8.2x10^-5^, one-tailed Wilcoxon signed-rank test), consistent with bisulfite-induced random fragmentation generating additional unique mapping positions beyond restriction enzyme cut sites. Within conversion methods, NRF increased with enzyme combination breadth (**Fig. 2d**, MH (0.31) > MT (0.24) > M (0.20)), with HaeIII showing the largest gain in capture site diversity.

#### eRREMS increases CpG recovery and maintains methylation reproducibility

At ≥5x depth per individual, eRREMS (enzymatic-converted MH) libraries captured 9.25 million CpGs, nearly twice the CpGs detected by standard RRBS (MspI bisulfite; 4.82 million; **Fig. 2e**), with consistent gains across coverage thresholds. The advantage was more pronounced when requiring coverage across all three individuals, with eRREMS capturing almost 3-fold more CpGs than standard RRBS at ≥10x depth (**Supplementary Fig. 2**), confirming that the expanded CpG set reflects reproducible capture rather than stochastic sampling. Pairwise Jaccard similarity indices showed that CpG sets clustered by library protocol rather than by individual (**Fig. 2f**), with eRREMS achieving the highest inter-individual reproducibility (mean Jaccard 0.71 vs 0.65 for MT).

Notably, enzymatic-converted libraries achieved greater read depth per CpG than bisulfite-converted counterparts across all enzyme combinations (**Fig. 2g**, Kolmogorov-Smirnov test, *P* < 2.2x10^-16^), with eRREMS libraries reaching a median depth of 7x versus 3x for standard RRBS. To decouple this advantage from differences in total sequencing yield, we performed random downsampling to matched read depths (**Fig. 2h**). Across all subsampling levels, enzymatic-converted libraries retained superior CpG recovery efficiency, with eRREMS showing the largest gains. At 37.5 million mapped paired-end reads, eRREMS recovered over 15 million CpGs; this depth was used for all subsequent analyses. Under matched read-depth conditions, methylation levels were highly concordant across conversion chemistries and enzyme combinations (mean Spearman’s ρ > 0.83; **Fig. 2i**), with samples grouping by individual within each conversion chemistry.

#### eRREMS redistributes CpG coverage toward distal, intronic, and CpG-poor regions

We next annotated CpGs by genic position and CpG island proximity in downsampled libraries (37.5 million reads, ≥5x per sample). Gains in CpG recovery with eRREMS were observed across all genomic categories (**Fig. 3a**) and were most pronounced in distal intergenic regions and introns. In distal regions alone, eRREMS captured over 3 million CpGs compared with ∼1.7 million in standard RRBS, with smaller but consistent gains in promoter-proximal regions and exons. Fold-change enrichment relative to RRBS across all library configurations is shown in **Supplementary Fig. 3a**. Stratification by CpG island proximity revealed a parallel pattern (**Fig. 3b**, **Supplementary Fig. 3b**). We observed the largest gains in shelf (2-4 kb) and open sea (>4 kb) regions, where eRREMS captured ∼4.5 million inter-island CpGs versus ∼2.3 million in standard RRBS. In contrast, enzymatic MspI did not show gain relative to RRBS in distal and open sea regions (**Supplementary Fig. 3a, b**), indicating that enzymatic conversion alone does not expand access to distal regulatory elements.

**Figure 3.**
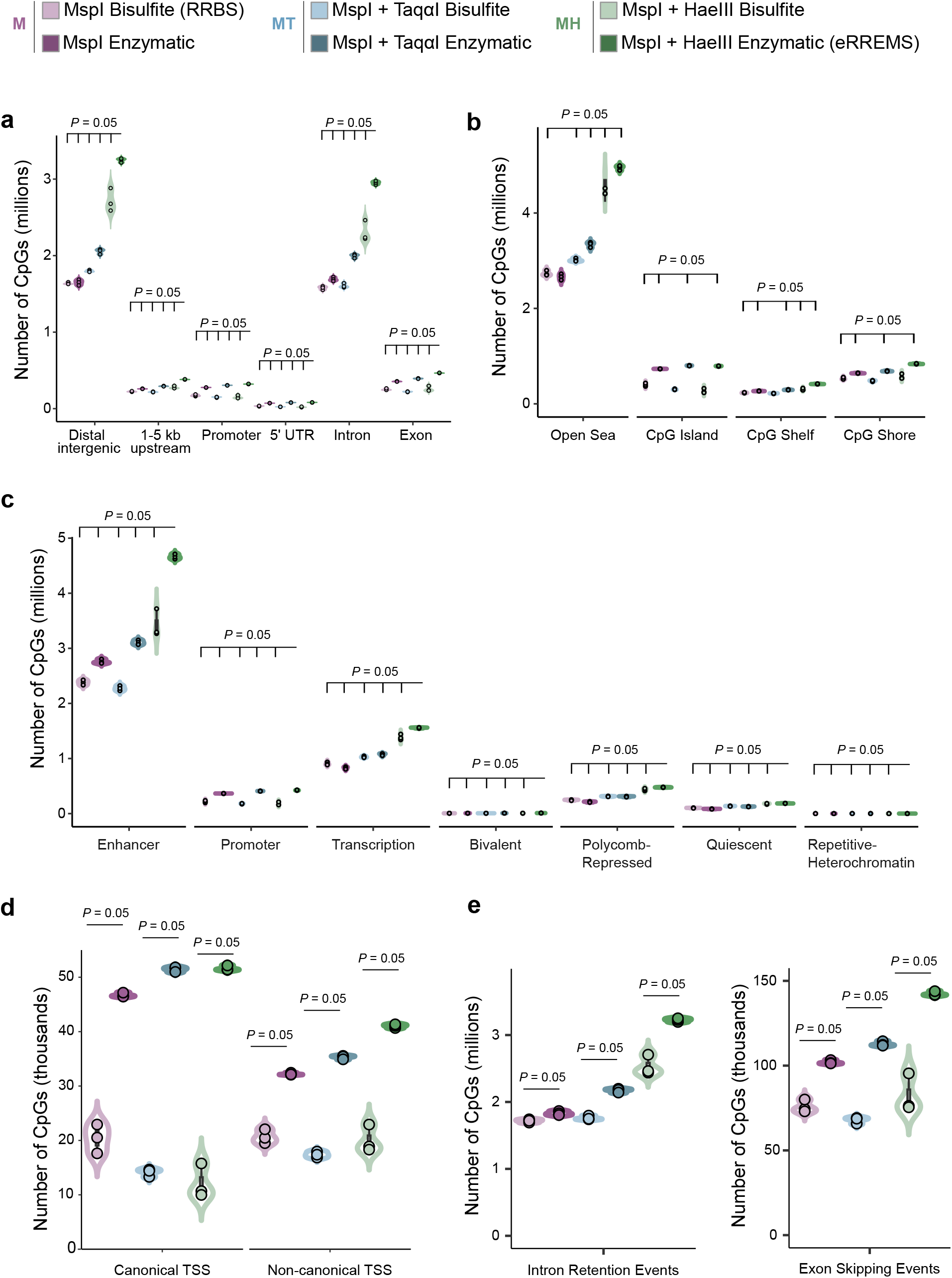
Genomic and regulatory annotation of CpG sites captured by eRREMS. All analyses were performed using downsampled datasets (37.5 million paired-end reads per library) considering CpGs detected at ≥5x coverage per sample **a.** Genic distribution of CpG sites detected across libraries. CpGs were classified according to their genomic position relative to annotated gene features (intergenic, promoter, 5′UTR, intron, exon). **b.** Distribution of CpG sites according to their proximity to CpG islands, classified as island, shore (≤2 kb from island), shelf (2–4 kb), or open sea (>4 kb from any island). **c.** Chromatin-state annotation of detected CpGs based on ChromHMM segmentations derived from 98 human tissues in the Roadmap Epigenomics dataset. Each CpG was assigned a single chromatin state (enhancer, promoter, transcription-associated, bivalent, Polycomb-repressed, quiescent, or repetitive/heterochromatin). **d.** Overlap between detected CpGs and transcription start sites (TSS) defined by CAGE data from the FANTOM5 project stratified by whether the TSS overlaps with an annotated gene promoter (canonical) or not (non-canonical). **e.** CpGs located within alternative splicing events according to VAST-DB annotations (left, intron retention; right, exon skipping). **a-e.** Significance was obtained with one-tailed Wilcoxon signed-rank test comparing MH enzymatic with each of the other libraries. Only significant comparisons (*P =* 0.05) are indicated. *P* = 0.05 represents the minimum achievable value for the sample size used (n=3 per group).

#### eRREMS extends methylome coverage into non-canonical regulatory elements

To determine whether expanded CpG recovery reflects access to functionally distinct regulatory elements, we annotated CpGs using chromatin state annotations derived from histone modification ChIP-seq profiles across 98 tissues and developmental stages from the Roadmap Epigenomics Project^30,31^. eRREMS captured >4.5 million enhancer-associated CpGs, approximately twice the number detected by standard RRBS (**Fig. 3c**), with 1.5 to 2-fold gains across promoter-associated, transcription-related, and Polycomb-repressed states (**Supplementary Fig. 3c**). TaqαI libraries showed disproportionate enrichment in repetitive and heterochromatin regions compared to RRBS, while enzymatic MspI libraries showed gains primarily in promoter-associated regions with negligible enrichment in enhancer-associated and other distal regulatory states (**Supplementary Fig. 3c**), suggesting that enzyme choice determines the functional composition of captured methylome. Orthogonal validation using FANTOM5 CAGE data on transcription initiation profiling across approximately 1,000 human primary cells, tissues and cell lines^32,33^ confirmed preferential detection of enhancer CpGs by eRREMS relative to standard RRBS (**Supplementary Fig. 3d**).

eRREMS libraries also detected substantially more CpGs at both canonical and non-canonical transcription start sites than standard RRBS (TSS, from FANTOM5^5,33^; **Fig. 3d**). Similarly, eRREMS showed extended CpG coverage at splicing-associated loci from VastDB (based on 1,478 RNA-seq datasets across diverse human tissues^34^; **Fig. 3e),** with more CpGs overlapping intron retention (3.2 million vs 1.8 million) and exon skipping (143,936 vs 74,364) events.

#### HaeIII uniquely captures dynamically regulated enhancers and non-redundant complex trait heritability

Having characterized the overall improvement in CpG recovery and library quality conferred by eRREMS, we next examined the functional relevance of the CpGs uniquely contributed by HaeIII. To isolate this contribution, we compared CpG overlap across restriction strategies. Enzymatic-converted libraries shared 3.8 million CpGs across all enzyme combinations (33.03% of total eRREMS CpGs; **Fig. 4a**), substantially exceeding the shared core in bisulfite libraries (2.8 million, 25.26%). Beyond this shared component, HaeIII contributed the largest enzyme-specific fraction: 4.2 million uniquely captured CpGs versus 1.5 million for TaqαI and 0.7 million for MspI.

**Figure 4.**
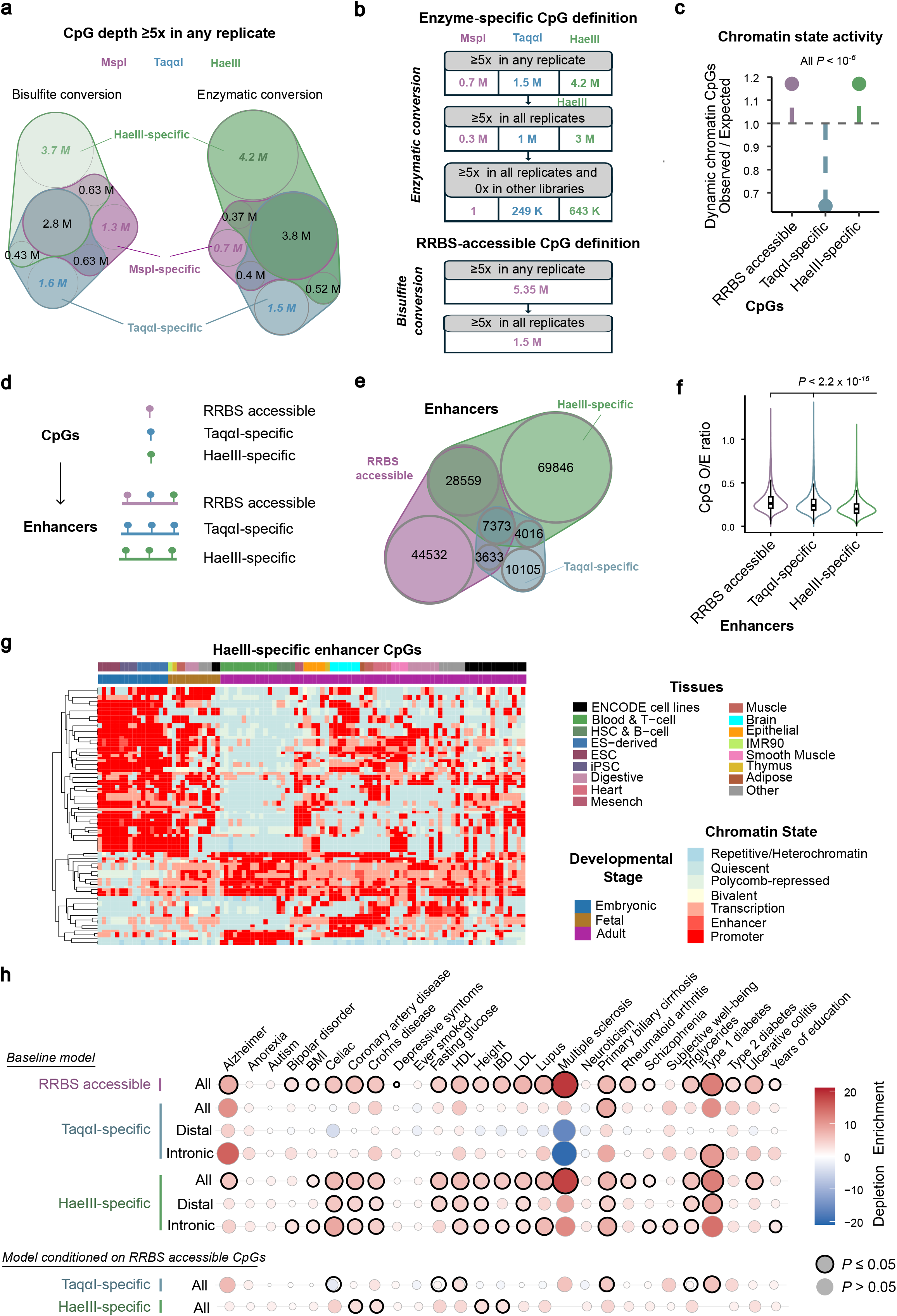
Biological relevance of regulatory elements uniquely accessed by eRREMS. **a.** Venn diagram showing the overlap of CpG sites detected at ≥5x coverage across restriction enzyme combinations for enzymatic and bisulfite libraries**. b.** Definition of high-confidence enzyme-specific CpG sets, requiring ≥5x in all three individuals of the library and zero counts across all replicates of other enzyme combinations. RRBS accessible CpGs were defined as those showing ≥5x in all three individuals in the MspI bisulfite-converted libraries. **c.** Observed versus expected ratio of dynamic CpGs (activity breadth 0.1-0.9) across CpG sets. P-values from Fisher’s exact test. **d.** Schematic illustrating the transition from CpG-level to enhancer-level definitions for each CpG category. **e.** Venn diagram showing the overlap between different enhancer classes. **f.** CpG O/E ratio for different enhancer classes. One enhancer RRBS with CpG O/E =3.847 was excluded from visualization. **g**. Chromatin state activity of HaeIII-specific CpGs located within HaeIII-specific enhancers across Roadmap Epigenomics reference epigenomes, ordered by developmental stage (embryonic, fetal and adult). CpGs were selected by filtering for activity breadth closest to 0.5 and subsequently ranked by significance of chromatin state transitions across developmental stages (Fisher’s exact test), resulting in 48 represented enhancers with 1-3 CpGs each. Each row represents one CpG and each column a tissue or cell type. **h**. Partitioned LD score regression enrichment coefficients for HaeIII-specific, TaqαI- and RRBS CpG sets, including distal and intronic subsets for enzyme-specific annotations, across a panel of complex trait GWAS. Upper rows show enrichment relative to the LD-MAF baseline model; lower rows show conditional enrichment after including RRBS accessible CpGs as an additional covariate. Circle size reflects the magnitude of the enrichment coefficient; circles with black outline indicate statistical significance (*P* ≤ 0.05). Red indicates positive enrichment; blue indicates depletion.

Applying a stringent criterion (≥5x depth in all three individuals and zero counts in all replicates of other enzyme combinations) yielded 643,017 HaeIII-specific and 249,365 TaqαI-specific CpGs in enzymatic-converted libraries (**Fig. 4b**). MspI-specific CpGs were effectively absent under these criteria (n=1), confirming library design integrity. Given that HaeIII-specific CpGs were predominantly located in intronic, distal intergenic and open sea regions (**Supplementary Fig. 4a, b**), we assessed their activity across the Roadmap Epigenomics reference epigenomes^30,31^. We defined dynamically regulated CpGs as those presenting active chromatin states across tissues and developmental stages (activity breadth 0.1-0.9 where 0 indicates inactive chromatin states across 98 epigenomes and 1 indicates constitutively active), reflecting context-dependent regulatory elements. HaeIII-specific CpGs were significantly enriched for dynamically regulated chromatin states (1.17-fold enrichment, Fisher’s exact *P* < 10^-6^), in comparable magnitude as RRBS accessible CpGs (≥1 CpG with ≥5x depth in all MspI bisulfite libraries, **Fig. 4b, c)**. In contrast, TaqαI-specific CpGs showed significant depletion for dynamic chromatin states.

We next asked whether HaeIII-specific CpGs define a repertoire of enhancer elements that is non-redundant with standard RRBS. Of the 109,794 enhancers containing HaeIII-specific CpGs, 69,846 (63.6%) were absent from standard RRBS libraries (**Fig. 4d, e**). The remaining 39,948 enhancers were also accessed by RRBS. In contrast, TaqαI-specific CpGs were associated with only 25,127 enhancers, the majority of which overlapped with the RRBS enhancer repertoire **(Fig. 4d, e)**.

HaeIII-specific enhancers showed lower CpG dinucleotide density relative to sequence composition (CpG observed/expected ratio, O/E) than RRBS-accessible and TaqαI-specific enhancers (**Fig. 4f**), consistent with their location in CpG-poor genomic regions. Together, eRREMS improves coverage within existing RRBS enhancers and additionally captures the CpG-poor but functionally dynamic HaeIII-specific enhancer space, expanding the total accessible enhancer repertoire 3.25-fold relative to standard RRBS (273,711 vs 84,097 enhancers), while retaining 99.5% of RRBS-accessible enhancers (**Supplementary Fig. 4c, d**). **Fig. 4g** shows representative HaeIII-specific examples of enhancers undergoing coordinated activation and silencing across embryonic, fetal and adult developmental stages, inaccessible to standard RRBS.

Because context-dependent regulatory regions frequently mediate disease susceptibility and phenotypic variability^35,36^, we applied stratified linkage disequilibrium score regression (S-LDSC)^7^ to determine whether genomic regions defined by HaeIII-specific CpGs are enriched for complex trait heritability. We found consistent and significant enrichment for these loci across 18 out of 27 complex traits, comparable to RRBS-accessible CpGs (**Fig. 4h**, top). This enrichment was reproduced when restricting analyses to only distal or intronic HaeIII-specific CpGs, and when RRBS-accessible CpGs were included as an additional covariate (**Fig. 4h**, bottom), demonstrating that HaeIII identifies regulatory regions contributing to complex trait heritability beyond those interrogated by standard RRBS. In contrast, genomic regions defined by TaqαI-specific CpGs showed no consistent heritability enrichment, either relative to baseline or beyond RRBS.

#### eRREMS libraries maintain stable CpG capture across preparations

To assess robustness of eRREMS, we compared the same three biological samples using independent implementations: two set of libraries prepared with the same enzymatic kit (Vazyme) and one prepared with an alternative enzymatic kit without physical size selection (NEB; Methods). CpG set concordance was highest between libraries prepared with the same kit and size selection strategy (mean Jaccard 0.727) and remained substantial when comparing libraries prepared with different kits and size selection conditions (mean Jaccard 0.696; **Supplementary Fig. 5a**). Across all protocol comparisons, within-individual concordance (mean Jaccard 0.706) exceeded between-individual concordance (mean Jaccard 0.578), indicating that biological differences between individuals contribute more to CpG set variation than preparation conditions. In contrast, fractional methylation estimates clustered strongly by individual (**Supplementary Fig. 5b**), confirming that eRREMS reliably captures biologically meaningful methylation differences.

#### eRREMS improves cohort-level CpG completeness at modest sequencing depth

WGBS distributes sequencing efforts genome-wide, resulting in a trade-off between breadth and per-site depth. In a deeply sequenced reference cohort of 205 methylomes across 39 cell types (>30x mean CpG depth^8^), WGBS detected 28.8 million CpGs with at least five reads in at least one sample, representing virtually all CpGs in the human genome (**Fig. 5a**). Imposing progressively stricter cohort-level coverage thresholds (≥5x in at least one individual, one third, two thirds, or all individuals) led to progressive loss of usable CpGs as per site read depth drops (**Fig. 5a**).

**Figure 5.**
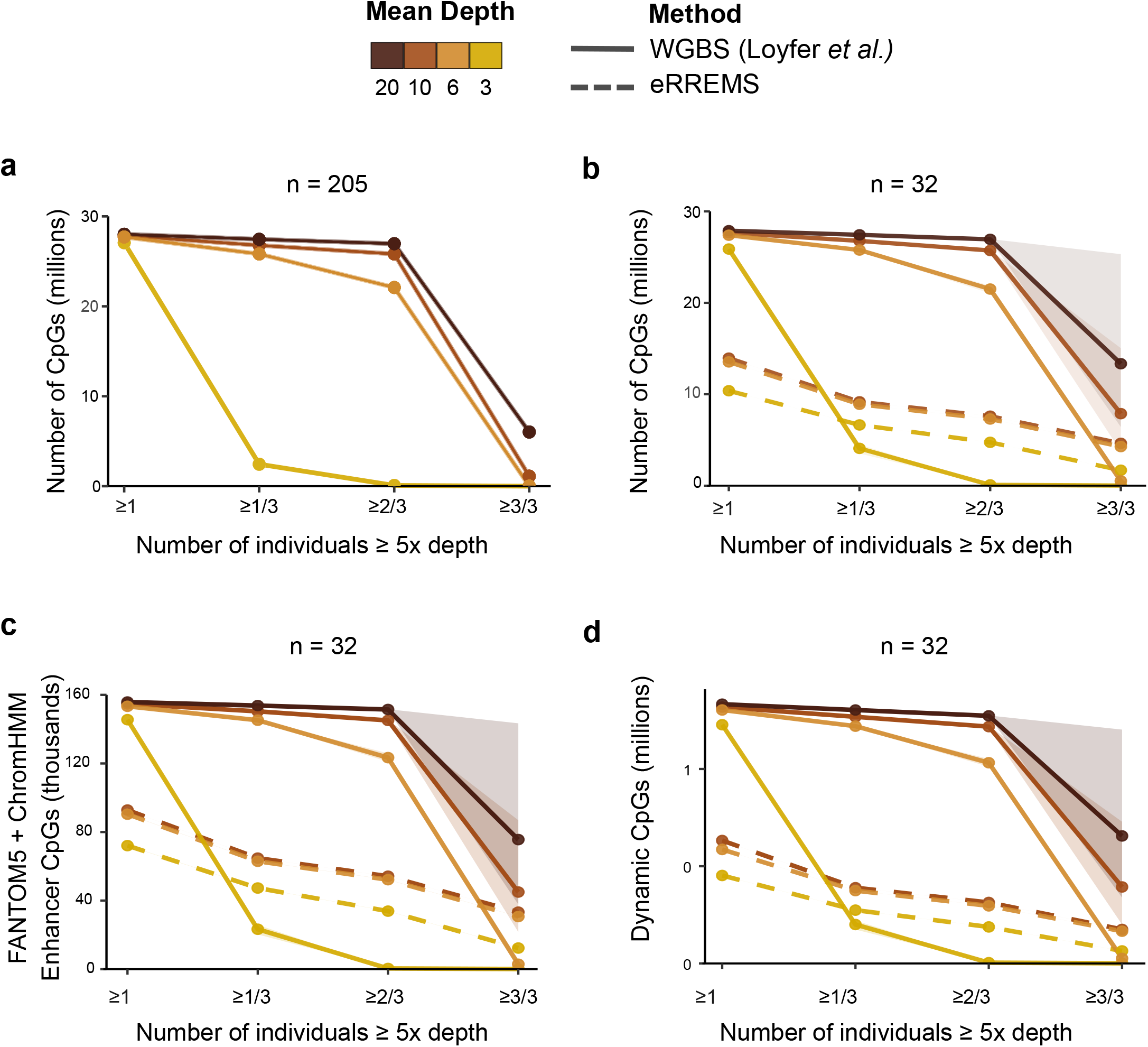
Cohort-level CpG coverage of eRREMS versus WGBS. **a.** Number of CpG sites detected in a cohort of 205 whole-genome bisulfite sequencing (WGBS) methylomes from the methylation atlas dataset (Loyfer et al). CpGs were counted if supported by ≥5x depth in at least one sample, one-third, two-thirds, or all samples in the cohort. **b**. CpG coverage across 32 sample subsets comparing WGBS and eRREMS libraries. **c**. Same analysis as in **b**, restricted to CpGs overlapping enhancer regions defined by ChromHMM annotations from the Roadmap Epigenomics dataset. **d**. Same analysis as in **b**, restricted to CpGs showing high variability in fractional methylation across the WGBS cohort (top 1% variance). **b-d.** WGBS curves represent random subsets of 32 samples drawn from the 205-sample cohort and downsampled to the indicated mean CpG depths. Shaded areas represent 95% confidence intervals (2.5^th^-97.5^th^ percentile) across ∼300 combined subsampling and binomial thinning iterations. Confidence intervals in panel b are too narrow to be visible, indicating high reproducibility across subsampling iterations

We expanded our eRREMS cohort to 32 samples and performed parallel subsampling analyses of the WGBS dataset to match sample size and per-CpG depth conditions (**Fig. 5b-d**; Methods). Although WGBS achieved broader total CpG coverage at high depth (20x), eRREMS CpG counts declined only modestly from ≥1 to ≥3/3 individuals across all depths examined, whereas WGBS showed progressive and substantial loss at each threshold. At 3x depth, eRREMS retained 1,690,563 CpGs consistently covered across all 32 individuals compared to 14,845 by WGBS (**Fig. 5b**). At 6x, eRREMS retained 4,275,494 CpGs versus 518,106 for WGBS. At 10x depth, this pattern was maintained, with eRREMS showing minimal decline across thresholds in contrast to continued WGBS loss at equivalent depth (**Fig. 5b**).

The completeness advantage was most pronounced for enhancer-associated CpGs. eRREMS retained 30,677 CpGs across all 32 individuals at 6x depth, whereas WGBS showed 2,877 at equivalent depth (**Fig. 5c**). At 10x depth, both methods retained comparable enhancer CpG numbers at the most stringent threshold (**Fig. 5c**). A parallel pattern was observed for variable CpGs (defined as the top 5% most variable CpGs by fractional methylation variation across the WGBS reference cohort; **Fig. 5d**). Together, these results demonstrate that eRREMS achieves greater cohort-level completeness across regulatory and dynamically variable CpG categories at substantially fewer reads than WGBS. In our data, WGBS required approximately 60% more paired-end reads than eRREMS to achieve comparable observed CpG depth (**Supplementary Table 4**). This estimate is likely conservative, as 150 bp paired-end reads generate overlapping sequences for a fraction of eRREMS library fragments (<300 bp; **Supplementary Fig. 1 a-c**); shorter reads of 100 bp would maintain equivalent CpG depth at reduced sequencing cost.

## Discussion

Reduced representation methylome profiling is constrained by two main limitations. Bisulfite treatment compromises DNA integrity and reduces reproducible CpG capture across samples, while MspI concentrates coverage in CpG-dense promoter-proximal regions. Here we show that both can be addressed simultaneously by substituting enzymatic conversion for bisulfite treatment and incorporating HaeIII as a complement to MspI. Bisulfite treatment shortened fragments, reduced sequencing yield, distorted base composition, and drove stochastic CpG loss independently of restriction enzyme choice, consistent with prior reports^25,26,29^.

HaeIII recognition does not require a CpG within the restriction site and generates complementary fragmentation patterns relative to MspI, increasing the likelihood that CpG-poor regions are recovered within size-selected sequencing libraries. Consequently, eRREMS captures a broader set of genomic contexts, including distal and intragenic regions where regulatory CpGs reside despite lower local CpG density. Enzymatic conversion further preserves fragment integrity, yielding longer inserts and greater per-CpG depth from the same sequencing input. Together, these two substitutions approximately double the recovery of regulatory CpGs relative to standard RRBS. Because sequencing cost dominates library preparation in most settings, incorporation of HaeIII as complementary enzyme substantially reduces the cost per regulatory CpG profiled.

CpGs gained by eRREMS are concentrated in dynamic enhancers, alternative promoters, and splicing-associated loci, as supported by multiple orthogonal experimental annotations. HaeIII addition expands the accessible enhancer repertoire 3.25-fold relative to standard RRBS, deepening CpG coverage within existing RRBS enhancers while accessing a substantially larger set of non-redundant regulatory elements. HaeIII-specific enhancers show lower CpG dinucleotide density than RRBS-accessible enhancers, consistent with GG|CC recognition being independent of CpG content and confirming access to a genuinely distinct regulatory lanscape. As CpG density has been shown to determine basal methylation states, with CpG-poor regions being preferentially subject to dynamic, context-dependent regulation^37,38^, the expansion of eRREMS into CpG-poor regulatory elements provides access to the most functionally informative fraction of the methylome. Partitioned LD score regression demonstrated that HaeIII-specific CpGs are significantly enriched for complex trait heritability, independently of the CpG space captured by standard RRBS, providing direct genetic evidence that the regulatory elements uniquely accessed by eRREMS harbor variants that are relevant for the genetic architecture of complex phenotypes.

TaqαI, the most widely adopted MspI complement in extended RRBS protocols^18^, showed disproportionate enrichment in repetitive and heterochromatic regions, depletion of dynamic enhancers, and no consistent heritability enrichment beyond standard RRBS, suggesting that TaqαI-based extended protocols may not substantially expand access to phenotypically relevant regulatory variation. Enzymatic conversion coupled with MspI has been recently proposed as an alternative to RRBS^24^. However, despite improvement in CpG recovery, functional gains were limited beyond promoter-proximal elements, indicating that HaeIII addition is the primary driver of eRREMS regulatory CpG expansion.

Although a decisive dimension of methylome profiling, cohort-level completeness is rarely considered among the primary criteria for method selection. As methylome profiling initiatives scale to ever larger sample sizes, the central challenge is not maximizing the number of theoretically accessible CpGs but ensuring reproducible quantification across individuals at sufficient depths for robust downstream analysis. WGBS maximizes breadth, but its genome-wide read distribution imposes an often-overlooked tension between per-site depth and cross-sample consistency. This trade-off is most acute for enhancer-associated and dynamically variable CpGs, which are increasingly recognized as central elements shaping cell fate and disease-associated regulation^39–41^. By contrast, concentrating sequencing efforts within a functionally enriched subset of CpGs, eRREMS achieves superior cohort-level completeness at modest depths with substantially fewer sequencing reads than WGBS. The stability of eRREMS coverage across cohort-level thresholds likely reflects the sequence-defined nature of restriction enzyme recognition, which targets the same genomic sites regardless of individual, in contrast to the stochastic read distribution of WGBS which becomes increasingly limiting as cohort size grows. Notably, eRREMS mitigates cross-sample missingness and depth heterogeneity, two factors that reduce statistical power in epigenome-wide association analyses^42^, and it preferentially profiles the set of regulatory CpGs that are most likely to yield discoveries in epigenome-wide association studies of complex traits.

The lower within-individual than between-individual CpG concordance across protocol implementations indicates that preparation conditions contribute less to CpG set variation than biological differences between samples, supporting reproducible methylome profiling across studies and laboratory settings.

Several limitations warrant consideration. We generated a total of 39 eRREMS libraries from fresh-frozen biological samples (including 3 benchmarking, 32 cohort expansion and 3 for alternative protocol application), sufficient for technical benchmarking but limited for assessing performance across diverse material conditions, including FFPE-derived, low-input, or degraded DNA commonly encountered in clinical and biobank settings. A direct comparison to enzymatic genome-wide methods such as EM-seq^23^ was not performed, as no suitable deeply sequenced cohorts were available. However, as these methods maintain genome-wide read distribution, similar constraints on per-CpG depth and cohort-level completeness are expected under equivalent sequencing budgets.

As epigenomic studies scale toward population-level cohorts^10,43^, sequencing-based multi-tissue atlases^6,8^, and single-cell DNA methylation atlases^44,45^, the limiting factor is increasingly not read depth per sample but the ability to recover functionally informative methylation signal consistently across the whole cohort at tractable cost. Rather than replacing comprehensive approaches, eRREMS addresses this bottleneck by offering a targeted and reproducible alternative for studies where regulatory depth and cohort-level consistency matter more than genome-wide breadth.

## Methods

### Sample processing and library preparation

Fresh frozen, OCT-embedded tissue specimens at prostatectomy were collected from localized prostate cancer patients at the Morales Meseguer University Hospital in Murcia, Spain. Samples were managed and provided by the Biobank IMIB (National Registry of Biobanks B. 0000859; PT23/00026), integrated into the Platform ISCIII Biomodels and Biobanks, following standard operating procedures with the appropriate approval of the Ethics and Scientific Committees (code CEIC-HMM 1/18).

DNA was extracted using standard protocol Qiagen AllPrep DNA/RNA Kit (QIAGEN, Cat. # 80204). 300ng of gDNA, 0.12 ng of Control DNA Unmethylated Lambda and 0.006 ng of control DNA CpG methylated pUC19 were digested with MspI enzyme (New England Biolabs, Cat# R0106T) and the same mix and amounts of DNAs were digested with TaqαI (New England Biolabs, Cat# R0149T) and HaeIII (New England Biolabs Cat# R0108T).

Prior to conversion, we performed a first double size selection with AMPure beads, using 0.6x and 1.6x bead ratios, followed by a second size selection using a 0.6x and 2.4x bead ratios, to enriched for DNA fragments between 100 and 500bp. Subsequently, 5ng of MspI digested DNA were combined with the same amount of MspI, TaqαI or HaeIII digested and size-selected DNA and this mixture was used for DNA conversion with “*EpiArt Magnetic DNA Methylation Bisulfite Kit*” (Vazyme, Cat. # EM103-01) or “*EpiArt DNA Enzymatic Methylation Kit*” (Vazyme, Cat. # EM301-01).

We prepared sequencing libraries using “*EpiArt DNA Methylation Library Kit for Illumina V3*” (Vazyme, Cat. # NE103-01) and “VAHTS Dual UMI UDI Adapters Set 1 for Illumina” (Vazyme, Cat. # N351), following version 22.1 of the kit manual. Briefly, after a DNA denaturation step, a truncated adapter was ligated to the converted ssDNA and an extension step was performed to achieve a complete dsDNA fragment. Following a clean-up step with SPRI beads, a second truncated adapter was added to the 5’ end of the original DNA strand. Next, after another clean-up step, ligation product was amplified through 14 cycles of PCR using index primers. The resulting library was then cleaned-up. See **Supplementary Note** for a detailed protocol.

Additional eRREMS libraries were prepared with NEBNext® Enzymatic Methyl-seq Kit (NEB Cat. # E7120S), following manualE7120 Version 7.0_4/23. In this case, 100 ng of MspI digested DNA were combined with the same amount of HaeIII digested DNA (no size selection was performed), and the mixture was used for library preparation.

#### Sequencing and quality control

Libraries were sequenced using pair-end 150bp reads with NovaSeq 6000. Sequenced library analyses were performed using standard pipelines. We performed a first quality control using fastQC (version v0.12.1, https://www.bioinformatics.babraham.ac.uk/projects/fastqc/). FASTQ files were adapter and quality trimmed using Trim Galore! (version 0.6.10, https://www.trimgalore.com/) with *trim_galore --illumina --clip_R2 12 --three_prime_clip_R1 12 –paired input1 input2 -o dir_outfiles*. Subsequently, we aligned trimmed reads to the human reference genome (NCBI hg38 primary assembly version 11) using Bismark aligner (Krueger & Andrews, 2011 (version 0.22.1) with standard parameters *-q --score-min L,0,-0.2 --ignore-quals --no-mixed --no-discordant --dovetail --maxins 500.* Methylation extraction was performed in two steps using Bismark tools. First, we used the extraction module to extract methylation status of cytosines, using parameters *-p, --no_overlap, --comprehensive*. Second, we generated the final cytosine reports with the merged methylation status of cytosines of both strands using *coverage2cytosine --genome_folder indexfolder --merge_CpG -o dir_outfiles*.

We calculated conversion and non-conversion rates using the *pUC19* and bacteriophage λ genomes (methylated and unmethylated controls, respectively). We extracted the conversion and non-conversion rates based on the methylated Cs in non-CpG (CH) and CpG context, respectively.

#### Insert size, base composition and complexity

We calculated insert sizes and base composition from mapped bam files using samtools: *samtools view S_XX_1_val_1_bismark_bt2_pe.bam | awk ’{print $9}’ > insert_sizes_SXX.txt* and *samtools view SXX_val_1_bismark_bt2_pe.bam | awk ’{read_id=$1; frag_len=$9; seq=$10; cg=gsub(/CG/, "", seq); print read_id"\t"frag_len"\t"cg}’ > SX_CG_content.txt*.

Library complexity was assessed using the non-redundant fraction (NRF), defined as the ratio of unique mapping positions to total mapped reads. Reads were filtered using samtools with the flags “-F 0x904 -q 30”, which excludes unmapped, secondary, and supplementary alignments and retains only reads with a mapping quality score ≥ 30. Total reads were counted across all retained alignments, and unique reads were counted after collapsing identical positional tags.

Pairwise Jaccard similarity indices were calculated between CpG sets detected at ≥5x coverage across all library pairs. The Jaccard index was defined as the number of CpGs covered in both libraries divided by the number covered in either library. Calculations were performed in R (version 4.4.1).

#### Read downsampling strategy

To allow comparisons between reduced representation libraries of different sizes, we followed a downsampling strategy. For reach individual library we randomly selected 13.5M, 25M and 37.5M reads from the bam files with Picard (https://broadinstitute.github.io/picard) as follows: *DownsampleSam -I input.bam -O output.bam -P 0.83945907 -STRATEGY Chained -RANDOM_SEED 42*.

#### Genic and CpG island annotations

CpGs were annotated using *annotatr R* package (version 1.32.0) and GENCODE gene information from UCSC hg38 annotation tracks. Because many CpGs overlap multiple features, we implemented a prioritization scheme to assign a single annotation per CpG. The scheme followed a two-step hierarchy: 1) feature type, ordered as protein-coding genes > non-coding genes > pseudogenes; and 2) genic context, ordered as promoter > promoter (5 kb) > 5′UTR > intron > exon > 3′UTR. Each CpG was assigned the annotation of highest priority according to this hierarchy.

CpG island context was annotated using the UCSC hg38 CpG island track (version 2022-10-1). CpG shores were defined as the 2 kb regions flanking islands, CpG shelves as the 2 kb regions flanking shores, and open sea as regions located >4 kb from any CpG island, following the AnnotationHub vignette (https://bioconductor.org/packages//release/bioc/vignettes/AnnotationHub/inst/doc/Ann otationHub.html).

#### Roadmap Epigenomics chromHMM annotations

We annotated CpGs using the chromHMM 18 chromatin states model derived from 98 epigenomes from the Roadmap Epigenomics Project (downloaded from https://egg2.wustl.edu/roadmap/web_portal/chr_state_learning.html). The chromatin states were grouped into seven functional categories: Promoter (TssA, TssFlnk, TssFlnkU, TssFlnkD), Transcription (Tx, TxWk), Enhancer (EnhG1, EnhG2, EnhA1, EnhA2), Repetitive/Het (ZNF/Rpts, Het), Bivalent (TssBiv, EnhBiv), Polycomb-repressed (ReprPC, ReprPCWk), and Quiescent (Quies). Because each CpG can overlap with multiple features across tissues, we applied a hierarchical prioritization scheme that assigns each CpG to its most regulatory-active state across tissues to avoid double counting. The scheme followed the following hierarchy in decreasing priority: 1) Enhancer, 2) Promoter, 3) Transcription, 4) Bivalent, 5) Polycomb-repressed, 6) Quiescent, 7) Repetitive-heterochromatin. This scheme emphasizes regulatory activity coverage of distal, low–CpG-density regulatory elements, which are typically underrepresented in conventional RRBS and are the features most likely to benefit from expanded restriction digestion. The same grouping and prioritization were applied identically to all libraries, so any bias introduced by this scheme is shared across protocols and does not affect relative comparisons.

To obtain a set of unique enhancers based on chromHMM data, we concatenated subsequent windows annotated as enhancer like state in any of the 98 tissues using the reduce function in GenomicRanges R package.

#### FANTOM5 annotations

FANTOM5-CAGE Peak annotations version 9 release (Jun 14, 2021) were downloaded from https://fantom.gsc.riken.jp/5/datafiles/reprocessed/hg38_latest/extra/CAGE_peaks/hg38_liftover+new_CAGE_peaks_phase1and2.bed.gz

#### VastDB annotations

CpG positions were annotated against the genomic coordinates of intron retention (INT) and exon skipping (EX) events retrieved from VastDB (Vertebrate Alternative Splicing and Transcription Database, hg38 assembly; EVENT_INFO-hg38.tab).

#### Dynamic enhancer definition and enrichment

Enhancers were defined as genomic regions assigned to enhancer-associated chromatin state (EnhG1, EnhG2, EnhA1, EnhA2 segments) in at least one of the 98 Roadmap Epigenomics epigenomes, as annotated by ChromHMM. Tissue breadth was calculated as the proportion of the epigenomes in which each enhancer was assigned one of these enhancer states. Dynamic enhancers were defined as those showing intermediate tissue breadth (0.1 ≤ breadth ≤ 0.9), reflecting elements that transition between active and inactive chromatin states across tissues and developmental stages, as opposed to constitutively active (breadth > 0.9) or constitutively inactive (breadth < 0.1) elements.

To assess whether HaeIII-specific CpGs are enriched for dynamic enhancer activity, we first identified enzyme-specific enhancers as those exclusively harboring HaeIII or Taq-specific CpGs. Then we performed a Fisher’s exact test comparing the proportion of dynamic versus non-dynamic enhancers between HaeIII-specific and non-HaeIII-specific enhancers sets.

For visualization of developmental dynamic enhancers, HaeIII-specific enhancers were first prioritized by having CpGs with dynamic breadth ∼0.5 and ordered by the p-value of a Fisher’s exact test comparing embryonic and fetal samples vs adult.

#### Enhancer CpG content

The ratio between observed and expected CpG dinucleotide frequency (CpG O/E) was calculated for each enhancer element using the following formula: CpG O/E = (N_CpG × L) / (N_C × N_G), where N_CpG is the number of observed CpG dinucleotides, L is the enhancer length in base pairs, N_C is the number of cytosines, and N_G is the number of guanines. Values below 1 indicate CpG depletion relative to sequence composition, as expected for most mammalian genomic regions due to methylation-driven CpG depletion over evolutionary time.

#### Partitioned heritability analysis using stratified LD score regression

We applied stratified linkage disequilibrium (LD) score regression^7^ to estimate the proportion of SNP heritability explained by CpGs detected by reduced representation enzymes. We constructed custom binary annotations for HaeIII-specific (n = 620,996 CpGs) Taqα-specific (n=123,090) and RRBS MspI accessible CpGs (n = 1,504,544). We expanded each CpG coordinate by 500 bp in each direction to capture SNPs within the region. A SNP was assigned a value of 1 if it fell within any expanded CpG region for a given enzyme, and 0 otherwise.

The baseline included^7,47^: (i) 10 minor allele frequency (MAF) bins (MAFbin1–10) to model MAF-dependent architecture; (ii) MAF-adjusted predicted allele age, which captures the relationship between allele age and per-SNP heritability under negative selection; and (iii) MAF-adjusted low LD scores computed from the African 1000 Genomes reference panel (MAF_Adj_LLD_AFR) which controls for background LD structure independent of population-specific demography. All continuous annotations were MAF-adjusted via quantile normalization within each MAF bin.

All files were downloaded from https://alkesgroup.broadinstitute.org/LDSCORE/. Partitioned LD scores were computed using LDSC with the 1000 Genomes Phase 3 European reference panel (--bfile 1000G.mac5eur), a 1 cM LD window (--ld-wind-cm 1), and restricted to HapMap3 SNPs (--print-snps). Regression weights and allele frequencies were obtained from the standard LDSC reference files (weights.hm3_noMHC and 1000G.mac5eur frequency files).

We performed two sets of analyses. First, we estimated the marginal heritability enrichment of each enzyme annotation separately (HaeIII-specific CpG versus baseline; RRBS MspI versus baseline, TaqαI-specific versus baseline). Second, we estimated the conditional heritability of HaeIII- and TaqαI -specific CpGs given RRBS Msp accessible CpGs by including both annotations in the same regression (HaeIII-specific versus RRBS MspI + baseline; and TaqαI-specific versys RRBS MspI + baseline), using the --overlap-annot flag. This directly tests whether HaeIII-detectable CpGs capture heritability beyond what is already explained by the standard RRBS enzyme. All analyses were performed using GWAS summary statistics with HapMap3 SNP filtering. Only autosomes were included and the MHC region (chr6: 25-34 Mb, hg38) was excluded.

#### Downsampling strategy for library size comparisons with WGBS

To enable fair comparison of cohort-level CpG completeness between eRREMS and WGBS under matched sequencing depth conditions, we implemented a CpG-level downsampling strategy based on binomial thinning of informative reads. This approach was applied to both eRREMS libraries and WGBS data downloaded from GEO (GSE186458). For cohort-level analyses, WGBS samples were randomly subsampled to match the cohort size of the eRREMS dataset (n = 32), and all downsampling procedures were repeated across 10 independent iterations to account for stochastic variation; results were summarized as means across iterations.

Downsampling targets were set to achieve observed effective CpG coverage depths of 3x and 6x for both eRREMS and WGBS libraries, with WGBS additionally downsampled to 10x and 20x observed depth to characterize performance across a broader coverage range. For eRREMS, analyses at ∼10x used natural sequencing depth (mean observed CpG depth 10.9x, range 9.6–12.7x); no downsampling was applied. For each sample, mean effective CpG depth was estimated as total informative coverage (sum of cytosine and thymine observations across all CpG sites) divided by the number of CpGs with at least one observation. A per-sample downsampling fraction was computed as the ratio of target to observed mean CpG depth, capped at 1 to avoid upsampling.

Binomial thinning was applied at the CpG level: for each CpG with observed coverage *k* ≥1, downsampled coverage was drawn as *k’ ∼ Binomial(k, f).* This preserves locus-specific depth heterogeneity while approximating reduced sequencing effort, avoiding the artificial inflation of cohort completeness that would result from uniform resampling across CpGs. After thinning, CpGs were classified as covered if *k’ ≥ 5*; cohort-level completeness was then summarized as the fraction of individuals meeting this threshold (≥1, ≥1/3, ≥2/3, or all individuals).

These analyses focus exclusively on CpG coverage metrics, which are independent of fractional methylation values and tissue-specific biology, allowing random subsampling across the diverse tissue types represented in the WGBS reference cohort without introducing biological bias.

#### Estimation of sequencing reads required for target CpG coverage depths

To estimate the number of paired-end reads required to achieve mean CpG read depths of 3x, 6x, 10x and 20x for eRREMS and WGBS, we applied a linear scaling approach based on empirically determined read-to-coverage relationships for each method. For eRREMS, read requirements were estimated from the relationship between total mapped paired-end reads and effective mean CpG coverage observed across our experimental libraries. For each target coverage depth *t*, the estimated read requirement was calculated as: *reads_needed = (median_reads × t) / median_coverage* where *median_reads* and *median_coverage* represent the median number of mapped pairedend reads and median effective mean CpG coverage across eRREMS libraries, respectively.

For WGBS, read requirements were estimated using publicly available sequencing metrics from Loyfer et al. 2023 (GEO accession GSE186458 and https://ega-archive.org/datasets/EGAD00001009789), without remapping of raw data. The median number of mapped paired-end reads, and median effective mean CpG coverage were calculated across the 205 samples in that dataset, and the same linear scaling formula was applied to estimate read requirements at each target coverage depth.

#### Variable methylation CpG definition

Dynamically variable CpGs were defined as the top CpGs ranked by variance in fractional methylation across the whole WGBS reference cohort (Loyfer et al., 2023). Fractional methylation at each CpG was calculated as the ratio of cytosine to total informative reads (C / C+T). Variance was computed across all samples with ≥1x coverage at that CpG, and CpGs were ranked accordingly. The top 5% most variable CpGs were retained for cohort-level completeness analyses.

## Availability of data and materials

The data generated in this project (raw and processed) are available in GEO and SRA (GSE341699). Code for data analyses are publicly available in Github at https://github.com/imendizabalCIC/eRREMS upon publication.

## Authorś contributions

U.L. performed the bioinformatic analyses, generated figures, and contributed to data interpretation and manuscript writing. M.G., N.M.-C. and L.B. performed the experimental work and library preparation. A.R., J.T., A.M., E.G.-B. contributed to sample collection, pathological and clinical annotation. A.C. contributed to conceptual discussions, project development, cohort access and co-supervised U.L. A.M.A. led the experimental design and supervised the experimental work. I.M. led the study, designed computational analyses, co-supervised U.L., coordinated the project, and wrote the manuscript. All authors discussed the results and approved the final manuscript.

## Acknowledgements

We thank Soojin V. Yi (University of California, Santa Barbara), Urko M. Marigorta (CIC bioGUNE) and Saioa García-Longarte (CIC bioGUNE) for critical reading of the manuscript and valuable feedback. We thank the Donostia International Physics Center (DIPC) for the use of high-performance computing (HPC) resources. We want to particularly acknowledge the patients and the Biobank IMIB (National Registry of Biobanks B.0000859, PT23/00026) for their collaboration.

## Supplementary Figure Legends

Supplementary Figure 1. **Pre-sequencing and sequencing output metrics. a.** Bioanalyzer electropherograms of each restriction enzyme digestion**. b.** Bioanalyzer electropherograms of each restriction enzyme digestion after size selection **c.** Bioanalyzer electropherograms of the final libraries per restriction enzyme digestion combination**. d**. Raw read counts after sequencing for all 18 libraries.

Supplementary Figure 2. Number of CpGs detected at each coverage threshold in all three replicates per library configuration.

Supplementary Figure 3. **Extended analysis of CpG distribution across library configurations. a-c:** Fold changes of annotation categories compared to standard RRBS (bisulfite-converted MspI libraries) as baseline **a.** Genic annotations. **b.** CpG island annotations **c.** ChromHMM annotations. **d.** Number of enhancer-associated CpGs across library configurations based on FANTOM5 CAGE annotations.

Supplementary Figure 4. **Genomic annotation and enhancer repertoire expansion by eRREMS**. **a.** Genic distribution of CpG sites detected across libraries. CpGs were classified according to their genomic position relative to annotated gene features (intergenic, promoter, 5′UTR, intron, exon). **b.** Distribution of CpG sites according to their proximity to CpG islands, classified as island, shore (≤2 kb from island), shelf (2-4 kb), or open sea (>4 kb from any island). **c**. Total enhancers detected by eRREMS, RRBS or both, defined as containing ≥1 CpG with ≥5x depth in all three individuals per library. **d**. CpG number detected by RRBS and eRREMS libraries for enhancers detected by both (n=83,715). Hexagonal binning was applied. Color reflects the number of enhancers per bin on a log scale. Dashed diagonal line indicates equal counts.

Supplementary Figure 5. **Robustness of eRREMS across library implementations. a.** Jaccard similarity heatmap showing the overlap of CpG sites detected at ≥5x depth across three independent library preparations of the same samples. Libraries include replicate digestions using the same protocol and libraries generated with an alternative enzymatic conversion kit and without physical size selection. **b.** Correlation of fractional methylation estimates at shared CpGs across independent eRREMS library preparations show in (a) from the same three biological samples.

## Supplementary Table Legends

**Supplementary Table 1. Fragment size distribution of *in silico* digestion products using independent or joint enzyme combinations.** Number of fragments generated by *in silico* digestion of the human genome with MspI, TaqαI, HaeIII restriction enzymes, either independently with fragments subsequently merged, or jointly considering all cut sites from both enzymes simultaneously. Fragments were classified into three size categories (<100 bp, 100-500 bp, and >500 bp) according to their length, with the total number of fragments generated per digestion strategy also reported. For reference, size selection in reduced-representation protocols typically target 100-500 bp fragments.

**Supplementary Table 2. Conversion rates based on unmethylated and methylated DNA controls.** Conversion efficiency was assessed using unmethylated lambda phage DNA and fully methylated pUC19 spike-ins included in each library. Conversion rates were quantified in unmethylated lambda at cytosines in CpG and non-CpG context. Over-conversion rates in pUC19 were quantified at cytosines in CpG context following standard practice. Each library is identified by a sample ID (S01–S18) indicating the library configuration, and an individual ID (P1–P3) indicating the biological donor.

**Supplementary Table 3. Mapping efficiency across all libraries.** Mapping statistics for all 18 libraries generated using the three restriction enzyme combinations (MspI, MspI + TaqαI, and MspI + HaeIII) and two conversion chemistries (bisulfite and enzymatic). For each library, the number of mapped read pairs and the corresponding mapping rate (%) are reported. Mapping rates were comparable across bisulfite- and enzymatic-converted libraries.

**Supplementary Table 4. Sequencing read requirements for eRREMS and WGBS at matched observed CpG depths.** Number of 150 bp paired-end sequencing reads required to achieve the indicated observed depths for eRREMS and WGBS datasets. Read counts correspond to the sequencing depth used for cohort-level completeness comparisons across depth thresholds in Fig. 5b-d.

## Supplementary Notes Legends

**Supplementary Note 1. Detailed experimental protocol for eRREMS and related benchmark library configurations.**

