## Supplementary figures and images for "eRREMS expands regulatory CpG coverage in reduced representation methylome sequencing"

### Supplementary Figure 1

##### Supplementary Figure 1

**a**

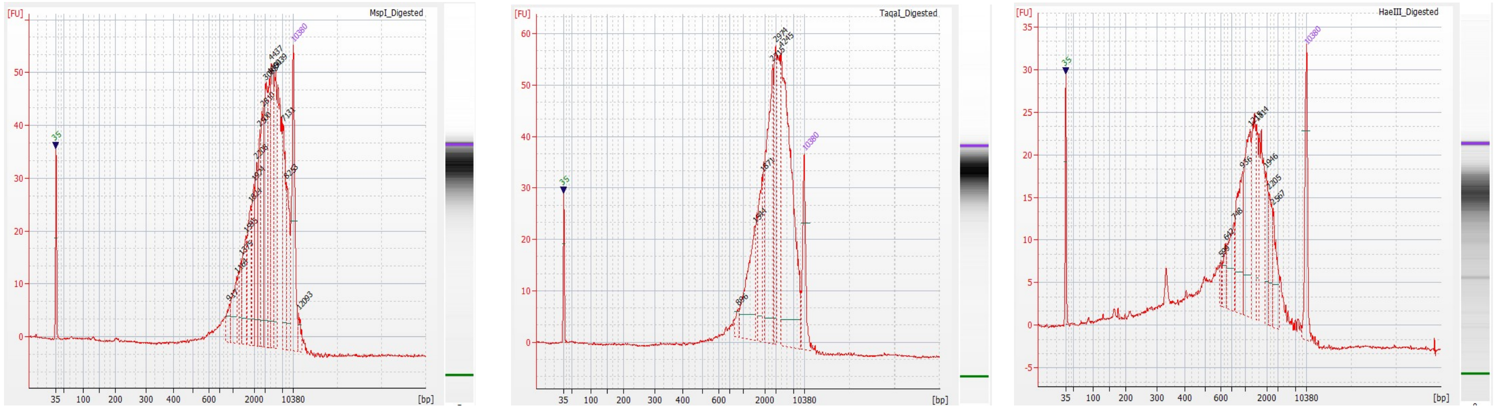


**b**

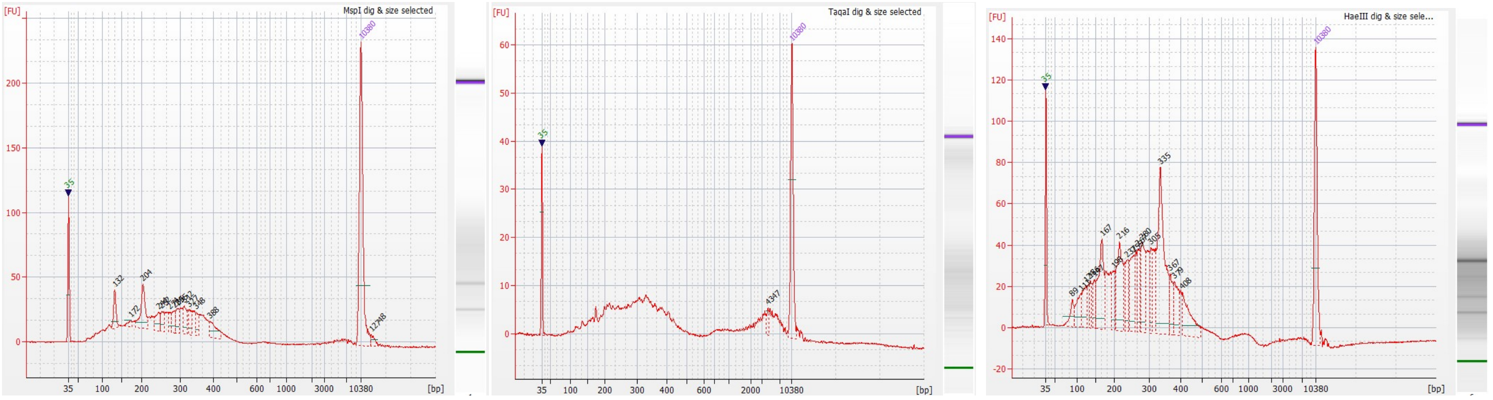


**c**

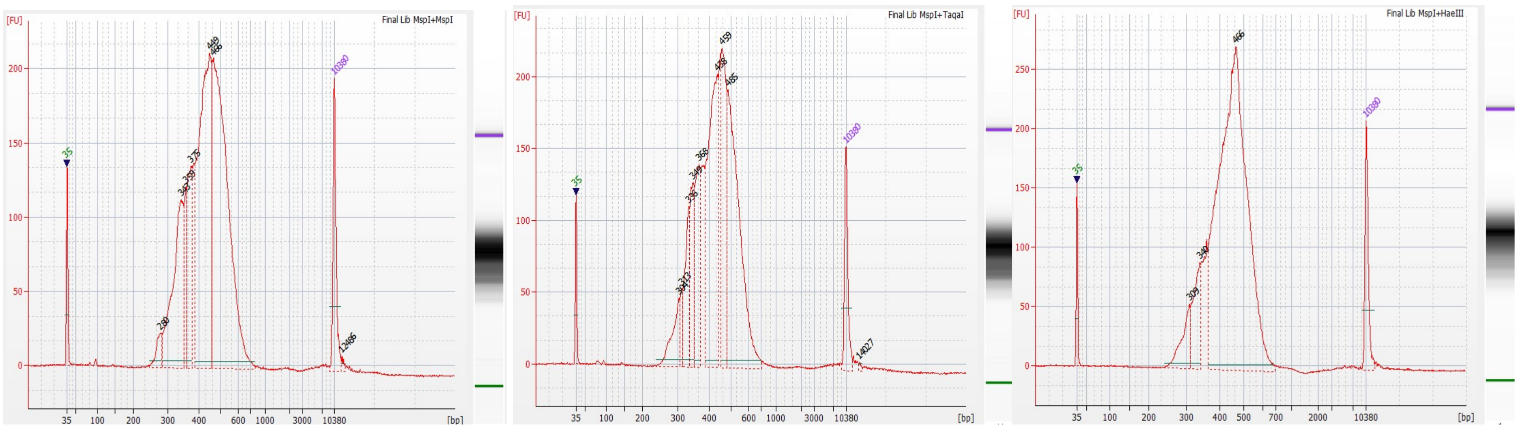


**d**

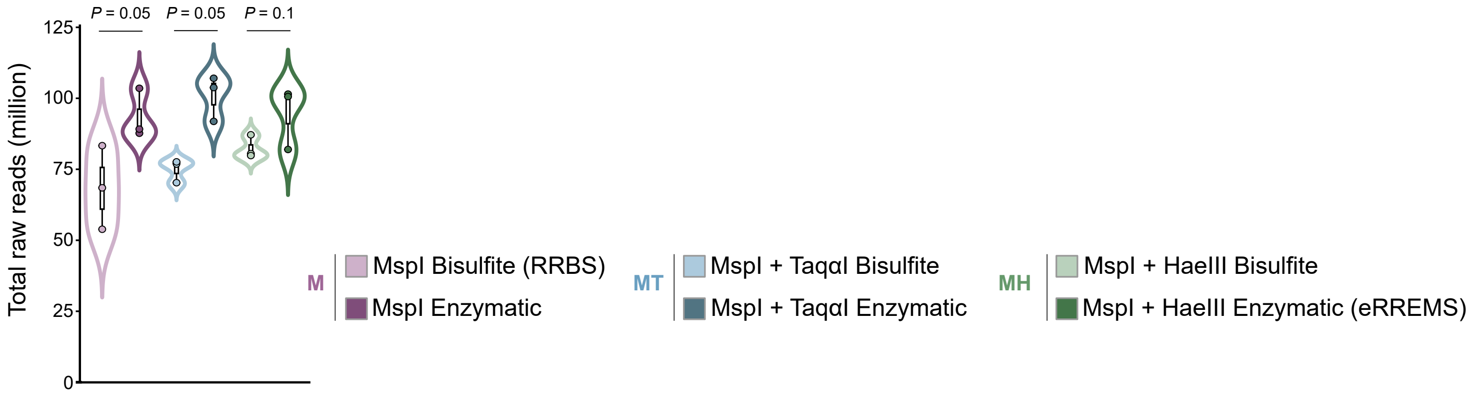

### Supplementary Figure 2

#### Supplementary Figure 2

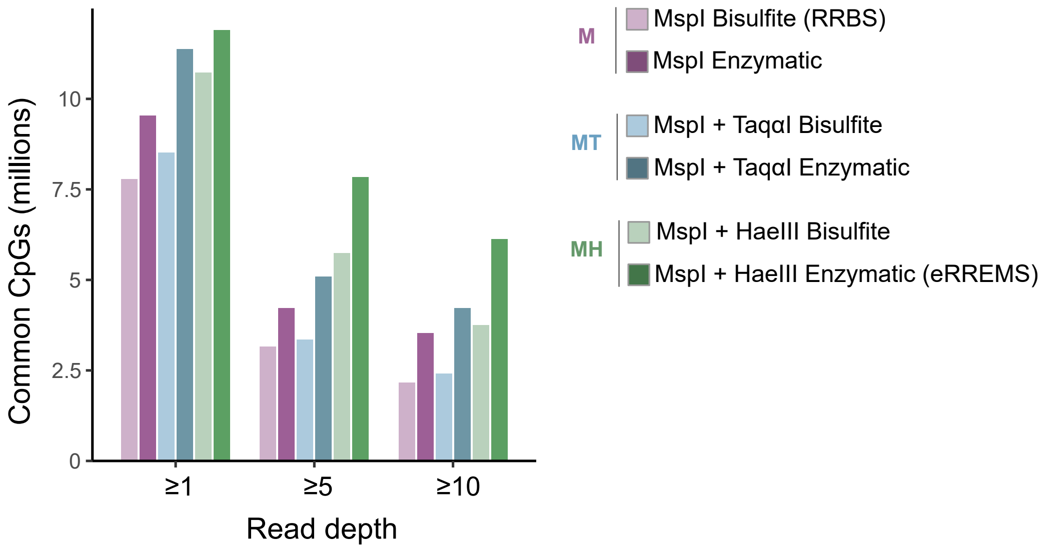

### Supplementary Figure 3

##### Supplementary Figure 3

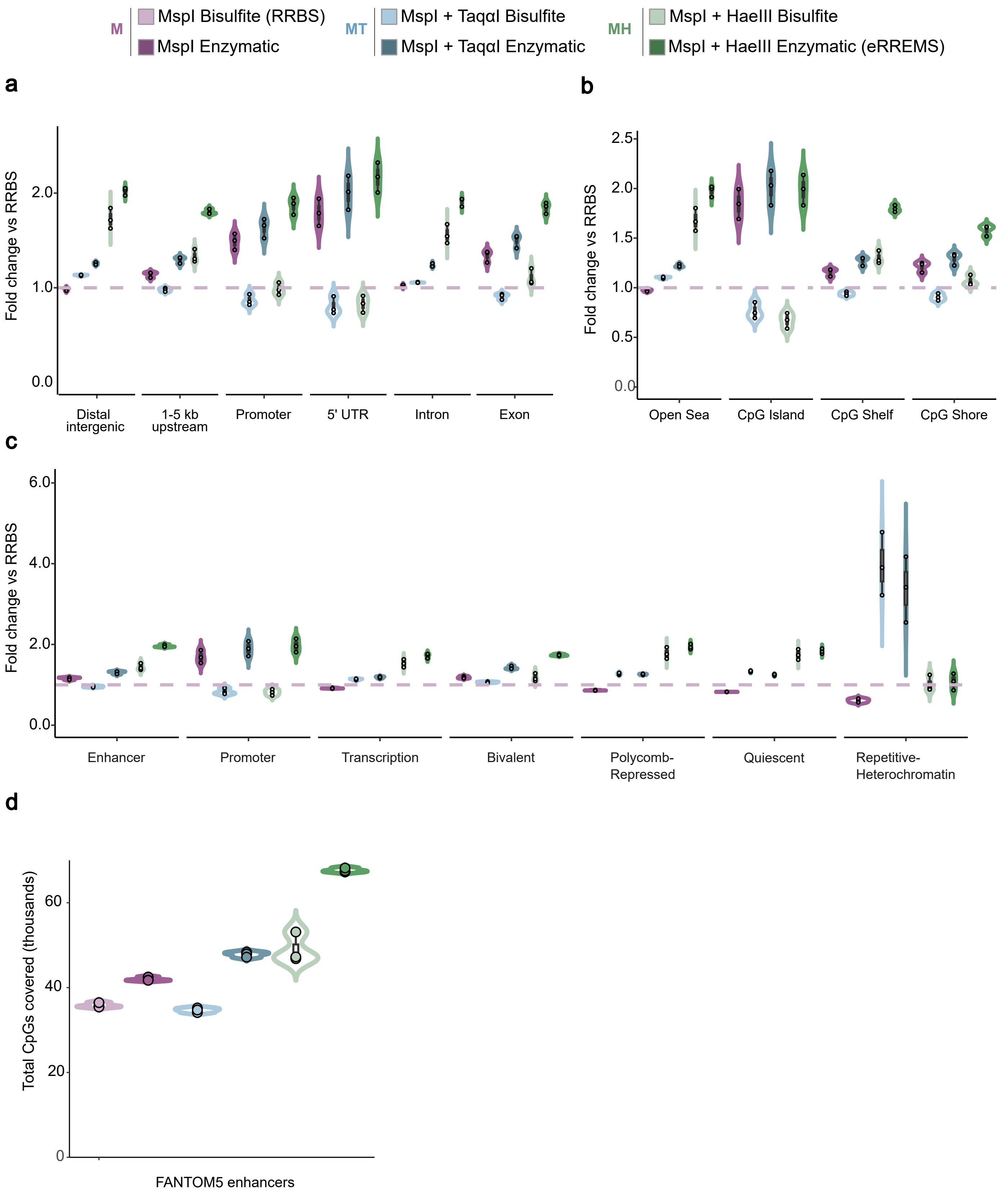

### Supplementary Figure 4

#### Supplementary Figure 4

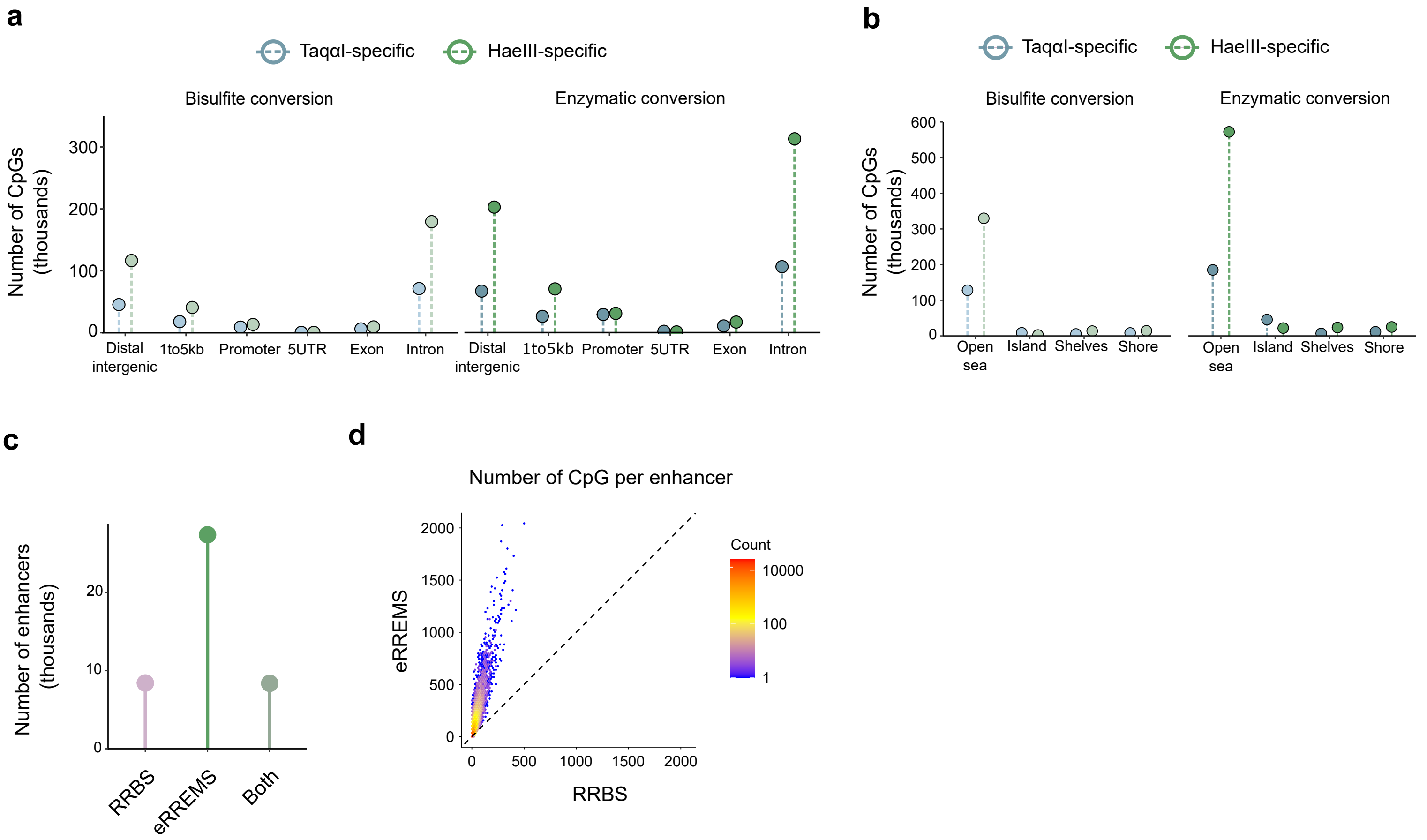

### Supplementary Figure 5

##### Supplementary Figure 5

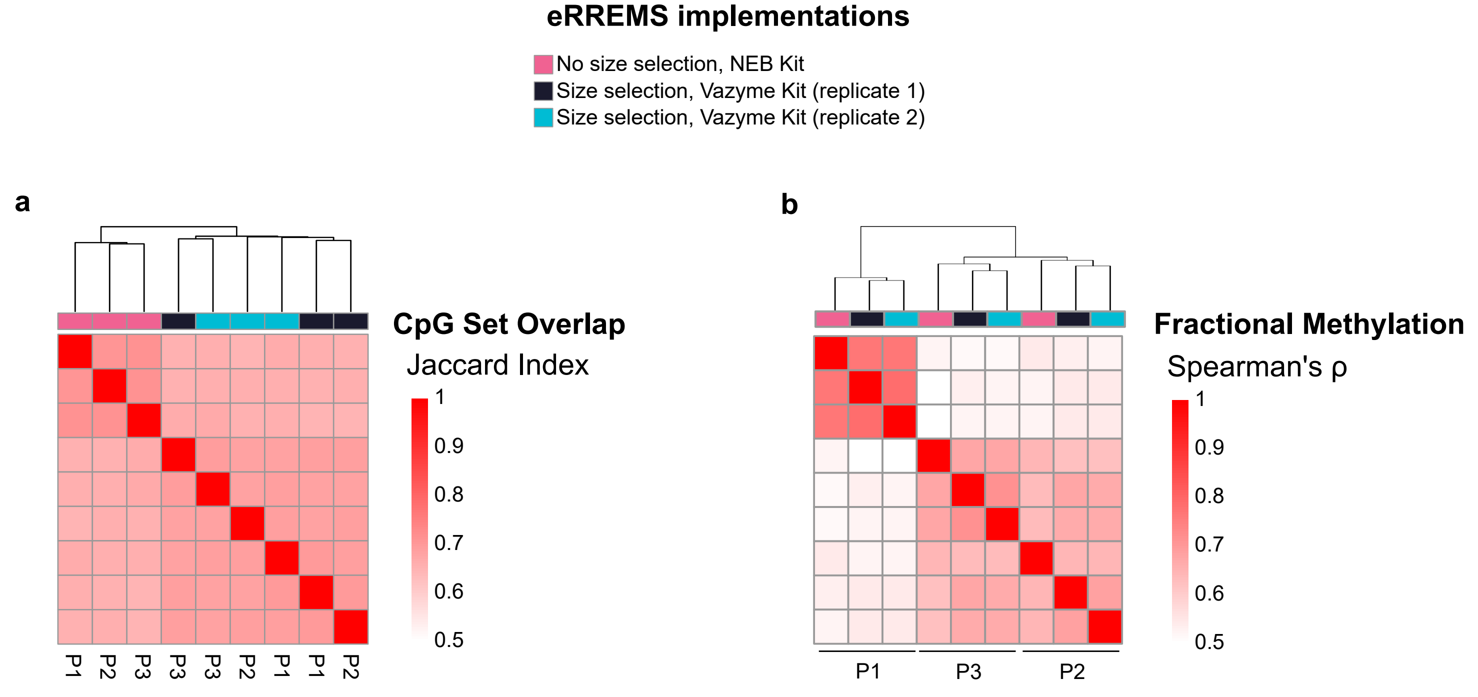
